# Computational constraints on the associative recall of spatial scenes

**DOI:** 10.1101/2022.10.08.511429

**Authors:** Kwang Il Ryom, Debora Stendardi, Elisa Ciaramelli, Alessandro Treves

## Abstract

We consider a model of associative storage and retrieval of compositional memories in an extended cortical network. Our model network is comprised of Potts units, which represent patches of cortex, interacting through long-range connections. The critical assumption is that a memory is composed of a limited number of items, each of which has a pre-established representation: storing a new memory only involves acquiring the connections, if novel, among the participating items. The model is shown to have a much lower storage capacity than when it stores simple unitary representations. It is also shown that an input from the hippocampus facilitates associative retrieval. When it is absent, it is advantageous to cue rare rather than frequent items. The implications of these results for emerging trends in empirical research are discussed.

## 1 Introduction

In one of the first recorded attempts to dissect declarative memory, Dante Alighieri, in *canto* XVII of his *Purgatorio* (*≈* 1315, see [1]), distinguishes 3 forms of increasing complexity of what he calls the *imaginative* faculty, the ability to recall facts and events not currently conveyed by sensory inputs. First, in verses 19-21, he exemplifies the ability to retrieve a single female character, from Greek mythology, from a cue comprised of a single major attribute – her metamorphosis from woman into nightingale; this can be accomplished, in principle, by a simple pattern associator [2]. Second, in twice as many verses, 25-30, Dante’s imagination recalls an entire scene, the death of Haman from the biblical Scroll of Esther. Although concisely reported, the scene is suggested to be more vivid and rich in detail than the paired associate recall, and also *compositional* in nature: it includes elements, like the death on the cross and the Persian Shah (in this instance, Ahasueros), which likely feature in several other unrelated scenes or events that the poet might have recalled. How can we conceive the neural mechanisms required to evoke such a compositional snapshot?

The third form of imagination, exemplified with an episode drawn from the poem by Vergil, Dante’s guide, is even more complex, since it involves Princess Lavinia speaking and trying to understand what went on in her mother’s mind that drove her to commit suicide. Recalling the episode therefore implies the recursive activation of theory-of-mind schemata: we need to get into Lavinia’s thoughts as she attempts to comprehend the dynamical unfolding of her mother’s drama by getting into her thoughts.

The ability to recall facts and events not currently conveyed by sensory inputs is integral to mind-wandering, the drifting of the mind away from current (sensory) experience towards inner contents such as memories or plans [3, 4]. Recent research has begun to investigate the neural underpinnings of mind-wandering, and to reveal distinct patterns of alteration of mind-wandering, following brain damage, which roughly parallel Dante’s poetic intuition of three levels of complexity in the imaginative faculty. Patients with lesions in the ventro-medial prefrontal cortex (vmPFC) tend to mind-wander less than healthy and brain-damaged controls, and when they do they are more focused on the present and on the self, suggesting a deficit in activating dynamical schemata to self-project into imaginary situations different from the perceptual present, such as future events or others’ perspectives [5]. Hippocampal patients, on the other hand, report mind-wandering as frequently as healthy controls, but their thoughts are of a streamlined logical/semantic character, impoverished in spatial details and bereft of episodic contributions, particularly from the recent past, the last year or so of actual experiences [6]. It thus appears that vmPFC integrity is necessary for the self-initiation and unfolding of mind-wandering episodes, whereas hippocampus integrity is important for the composition of elements drawn from recent experience into imagined scenes that fuel mind-wandering, whether or not they closely match combinations of elements that actually occurred [7, 8, 6, 9, 4].

Why should it be so? After all, influential memory theories promote the idea that, after hippocampally-driven consolidation, even episodic memories should become independent of the hippocampus [10, 11]. One such theory viewed the hippocampus as a complementary learning system, needed because the cortex, just like a back-propagation trained network, is postulated to be able to only learn slowly [12]; logically, once the cortex has taken its time, the hippocampus can be disposed of. The Multiple Trace Theory has emphasized instead the qualitative distinction between truly episodic memories that remain dependent on the hippocampus through a lifetime, and semanticized memory content that can be retrieved and utilized also without the hippocampus [13]. A somewhat intermediate formulation has been put forward, to try and reconcile the contradictory empirical evidence, which can be invoked in partial support of either extreme position: it holds that the hippocampus regenerates constructs that appear to be simple reactivations of the activity patterns originally encoded, but are not [8]. By titrating the degree of infidelity of the reactivated from the original, this proposal can satisfactorily interpolate between views that *prima facie* clash with each other.

None of the above, however, really addresses any constraints that may arise below the functional system level, that is, in the neural network mechanisms that are invoked to implement the required operations of memory storage and reactivation. An exception may be the argument that rapid neocortical learning would lead to catastrophic interference [12], although it was later qualified that this would only happen with new content *inconsistent* with previously stored information [14]. Episodic memories, however, are typically neither fully consistent nor inconsistent with each other, rather, they are diverse, entailing a variably overlapping set of items.

We ask here whether there are purely computational constraints that require cooperation between the hippocampus and neocortex in the associative storage and retrieval of snapshot compositional memories, and which stem from the distinct neural network organization of the hippocampus and of vmPFC (and neocortex in general). The hippocampus has available the dentate gyrus, which can establish a new, tendentially orthogonal compressed representation for any new memory [15]. In the neocortex there is no dentate gyrus, but its presumably large storage capacity – particularly in humans – should allow for the associative storage of many new combinations of items, most of which are already endowed with their neuronal representations. To what extent is this the case?

## 2 The Potts model of an extended memory network

The Potts associative network offers a tractable mathematical model (as shown from the early study by Ido Kanter [16]) of how compositional memories may be stored over large swathes of neocortex and later retrieved through purely associative mechanisms. In contrast to the neural networks usually considered in machine learning, it is recurrent (also at large scale) rather than directional, lacking *a priori* defined input and output layers, and the connection weights are modified by a model associative rule, with no involvement of error signals. As a memory device, therefore, it operates by self-organization, without instructions or extra ingredients, as envisaged by Braitenberg [17]. Its core idea is that it focuses on the interactions between patches of neocortex, mediated by axons travelling through the white matter, subsuming instead the dynamics within each patch into the suitably defined Potts units. The correspondence with the underlying full-fledged cortical network has ben discussed in [18].

### 2.1 Potts units that model cortical patches

A Potts unit, introduced in statistical physics in 1952 [19], has *S* states pointing each along a different dimension and can be regarded for our purposes as representing a local subnetwork or cortical patch of real neurons, endowed with its set of dynamical attractors; these span different directions in activity space, and are assimilated to the states of the Potts unit. One can then define an autoassociative network of *N* Potts units interacting through tensor connections [18]. The memories are stored in the weight matrix of the network during a learning phase, as assumed in the Hopfield model [20]: each memory *µ* is a vector or list of the states taken in the overall activity configuration by each unit 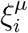. We label by the index *k* each of the *S* states a Potts unit can take (assuming for simplicity *S* to be the same throughout the network) and we consider an extra quiescent state, *k* = 0, when the unit does not participate in the activity configuration of the memory. Therefore *k* = 0, …, *S*, and each 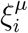 can take values in the same categorical set.

For the standard, quasi-orthogonal memories, for which it is assumed that units are assigned an activity state independently of each other and of any information already present in the network, the tensor weights read [16]

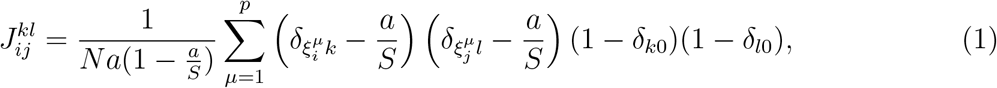

where *i, j* denote units, *k, l* denote states, *a* is the fraction of units active in each memory, and the *δ*’s are Kronecker symbols. The subtraction of the mean activity per state *a/S* ensures a higher storage capacity [21]. We shall see how this *learning rule* has to be modified if we consider, in the case of compositional memories, that information about their components is already present in the network.

Locally, each patch of cortex tends to align its activity with one of the attractor states or to remain in the null state, which is expressed in the Potts model by

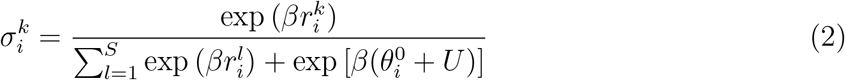

and

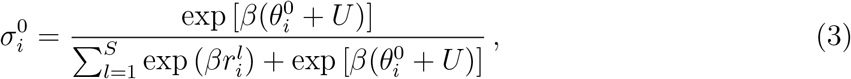

where 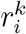is the input variable tending to align unit *i* to state *k*. This occurs within a time scale *τ*_1_, as expressed by the equations below. *U* and 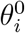 are the common and unit-specific components of an effective threshold, that incorporates a variety of inhibitory effects. From Eqs. (2) and (3), we see that 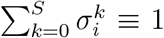, and note also that 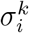 takes *continuous* values in the (0,1) range for each *k*, whereas the memories, for simplicity, are assumed discrete, implying that perfect retrieval is approached when 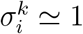 for 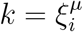 and *≃* 0 otherwise.

### 2.2 Potts model dynamics

When the Potts model is studied as a model of cortical *dynamics, U* (*t*) and 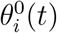 are functions of time, to represent the temporal course of long-range (indirect) and local inhibition. In addition, state-specific thresholds 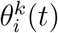 are used to represent firing rate adaptation effects [22]. Here, however, we are only interested in modeling associative retrieval of a content-addressable memory, that is, a simple rapid operation that succeeds or fails within the time scale *τ*_1_ of neuronal activation. We do not consider, therefore, slow inhibition and adaptation processes, and assume for simplicity a single common component of the threshold, *U* (*t*), which varies rapidly enough to regulate the overall activity in the network at a constant level,

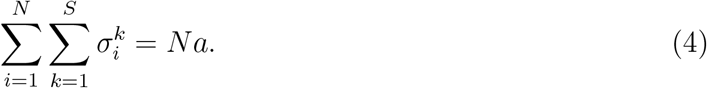

While this assumption is rather unrealistic, it allows assessing the associative capability of the network under ideal conditions, free of the temporal fluctuations of unregulated dynamics. The time evolution of the network is then governed by the equations

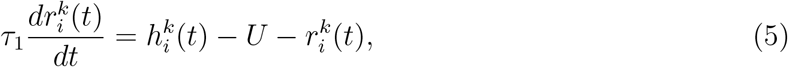

where *U* is adjusted instantaneously to maintain Eq. (4). The input that the unit *i* in state *k* receives is the sum of an *internal* component, i.e., coming from the other Potts units, and a term 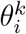, which here is used to model not adaptation but a sustained external signal, considered constant over the time scale of retrieval, such as that which can come from the hippocampus (we use it with a minus sign for consistency, as *θ* usually denotes a threshold).

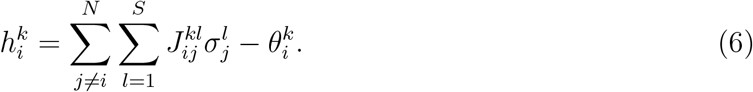

The way the patterns 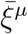 are generated, i.e. their probability distribution, has effects on the retrieval properties of the network, i.e. on its ability to retrieve with good accuracy one of the stored memories, if this is partially cued. A quantitative measure of this ability of the network is the *storage capacity*, the number of memories the network model is able to store and retrieve, given values for its various parameters.

### 2.3 Simple memories

If memory patterns are nearly orthogonal, that is, randomly correlated, like those assumed to be established by the dentate gyrus in the hippocampus, the Potts model equipped with Eq. (1) can store and retrieve an extensive number of patterns and each pattern has a large basin of attraction (Fig. 2a,b).

What if memory representations have a nontrivial structure, rather than randomly correlated? In the next section, we examine the retrieval properties of Potts neural networks when memories have a semi-naturalistic internal structure, in terms of items.

**Table 1.** Parameters of the network

| Symbol | Meaning | Default value |
| --- | --- | --- |
| $N$ | number of Potts units | 1000 |
| $S$ | number of states per unit | 7 |
| $p$ | number of memory patterns | 200 |
| $a$ | sparsity of patterns | 0.2 |
| $Z$ | number of items per memory | 5 |
| $K$ | number of items in total | 200 |
| $B$ | number of bins | 20 |
| $f$ | fraction of units for cuing | 0.5 |
| $\gamma$ | strength of hippocampal input | 0.1 |

### 2.4 Compositional memories

We take our model compositional memories to include *Z* items, each of which has a distributed cortical representation, like simple memories in the hippocampus (Fig. 1b). Across memories, some items may appear more frequently than others. We consider a pool of *K* items. Each memory can contain items with different frequency, from rare to very frequent ones.

**Figure 1.**
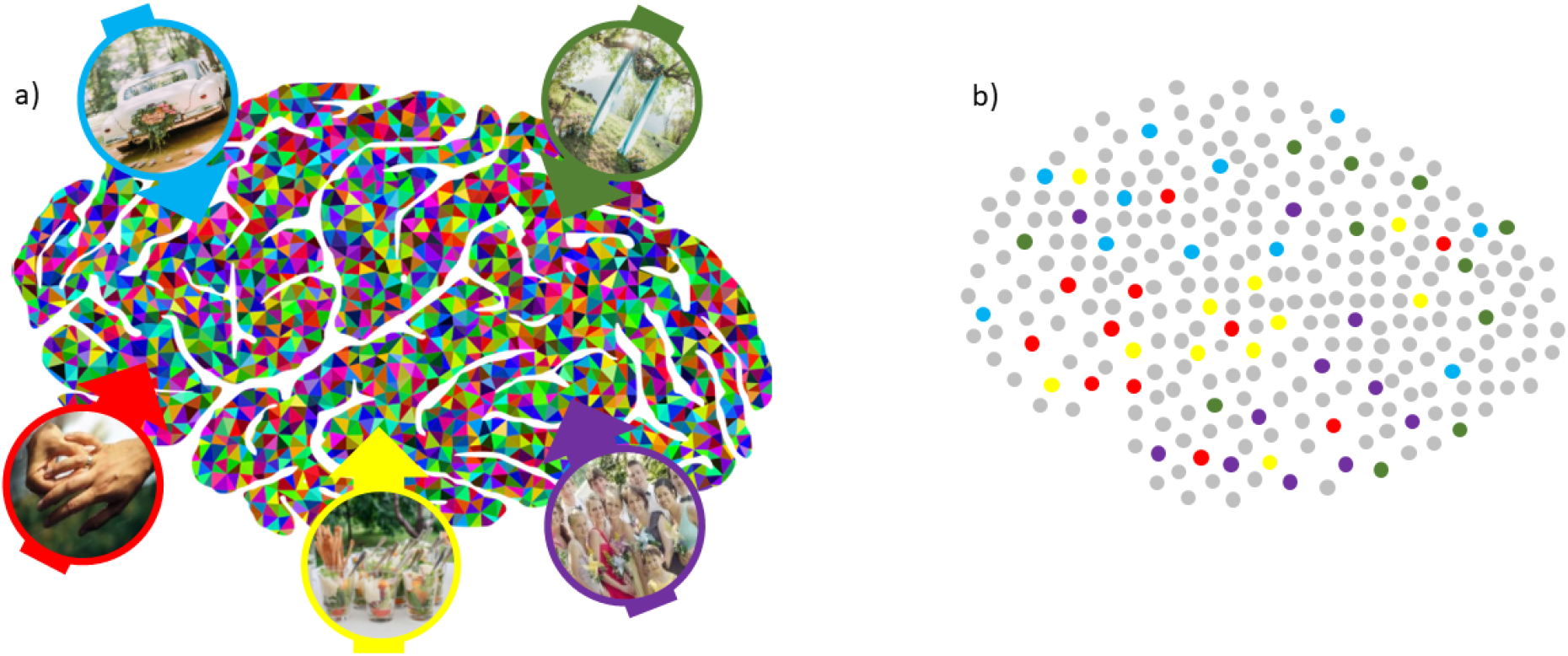
Compositional memories as representated in a Potts neural network. The episodic memory of a friends’ wedding relies on a composition of items already given, at least in some instantiation, a long-term representation **(a)**, such as the canopy, the people present, the food, the rings and the car with which they leave. In the Potts model **(b)** these elements are assigned sparse distributed representations over several Potts units, each loosely standing for a small patch of cortex.

We denote with 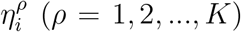, the activity patterns, of sparsity of *a*^*′*^ = *a/Z*, which represent the items. Here *a* is the sparsity, i.e., the fraction of active units, of the memories themselves, 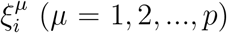, and the details of how we compose the memory patterns from those of the items are explained in the Appendix.

The connection weights are set differently than in Eq. (1), to express the notion, inherent to the compositional construct, that once an item has been encoded onto the synaptic connections, it is there and it is not stored again every time that item is present in the input

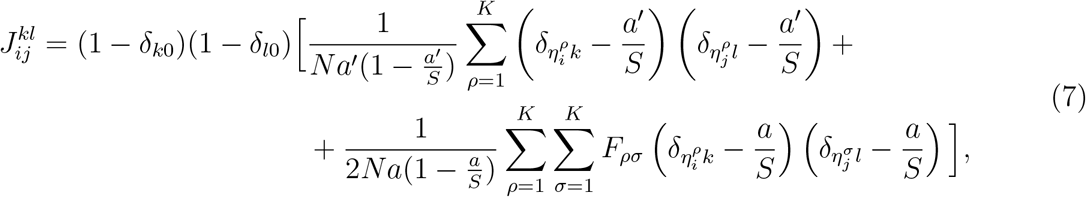

where *F*_*ρσ*_ is 1 if a pair of items (*ρ, σ*) appears together in one of the *p* memories, and 0 otherwise. That is, while the first term of Eq. (7) reflects one-shot associative learning of individual items, assumed to have occurred before, the second term likewise stores relations between items included at least once in the same compositional, episodic memory, and again the pair is stored once even if it recurs in multiple memories. Note that the prefactor with *a*^*′*^ in the denominator makes the single-item term stronger than the pair-of-items term, as 1*/a <* 1*/a*^*′*^ = *Z/a*. Note also that more complex, e.g., iterative and non-associative processes involved in acquiring the individual items in memory are not considered in the present model for simplicity, but they would not necessarily affect the constraints we focus on here, which are those arising from the associative storage not of items but of unique compositions of items.

### 2.5 Retrieval cues *vs*. Hippocampal input

To simply *cue* the network we activate a fraction *f* of the units active in a given memory, concentrated within some of the items of that memory, and let the network evolve without further external input. For example, for *f* = 0.5, when *Z* = 5, the cue is applied essentially to all the units active in two of the items, and to half of those active in a third (minor adjustments are due to the coincidence of some of the active units). With memories including both rare and frequent items, we consider applying a cue concentrated at either end of the frequency spectrum.

To model hippocampal inputs operating at retrieval, instead, we assume that the hippocampus has reinstated a compressed representation of the entire memory, and is able to convey a corresponding signal to all the units of the Potts network, which unlike the cue is sustained over the time course of retrieval. We express that through the state-specific thresholds, 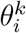, by setting, for memory *µ*,

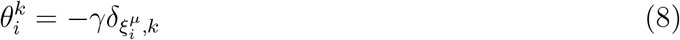

so that *γ* regulates the intensity of facilitation. Note that this 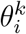is taken to be constant in time.

## 3 Results

### A strong constraint on compositional memories

First, for the sake of analytical clarity, we start from a simple case, in which all items appear with the same (average) frequency in the compositional memories: we vary the number of memories *p*, compose each by drawing from a common pool of *K* items, and set the other parameters at their default values, specified in Table 1, including the number of items per memory *Z* = 5. Note that, when for example *p* = 300, items appear on average in 5 distinct memories each, if *K* = 300 as well, and in as many as 15 memories each, if *K* = 100. This increases the difficulty of maintaining the unique item configuration of the compositional memory, even though it is present in the full cue (Fig. 2a), and once *p* = 400, compositional memories are effectively inaccessible (the overlap, which measures the correlation of the retrieved activity with the stored representation, drops to zero); whereas simple unitary memories (which can be conceived as comprised of non-repeated items) do not show a capacity limit, with our parameters, until *p* = 16000.

**Figure 2.**
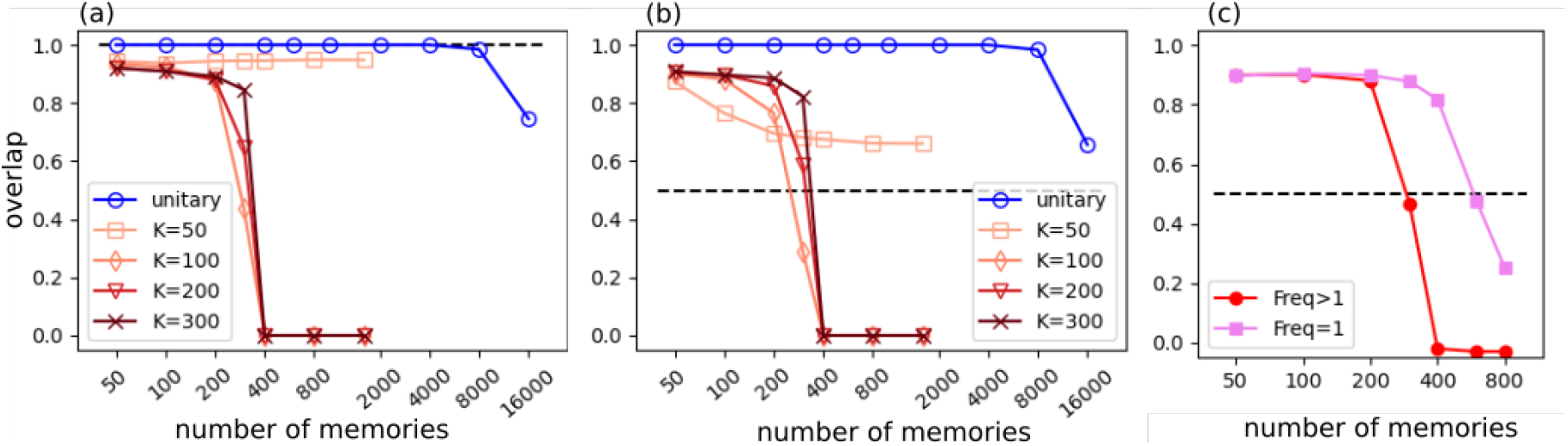
Unitary memories and compositional memories. **(a)**: The overlap between stored patterns and retrieval states is plotted as a function of the total number *p* of stored memories. The network stores only one type of memory: either unitary (blue) or compositional memories with fixed frequency (red). The network starts from a perfect version of one memory pattern (*f* = 1.0) and is allowed to follow its dynamics until it reaches a retrieval state or a limiting time. From bright red to dark red, colour encodes the number of items *K*. The blue curve is for random patterns (unitary memories). Network parameters are set at default values (see Table 1). **(b)**: Similar to (a), but the network is partially cued by a memory pattern. The partial cue is prepared by flipping back a fraction 1 − *f* = 0.5 of active units of the cued pattern into a quiescent state. **(c)** The same network stores two types of compositional memories: memories made of frequently used items (red) and those made of rarely used items (used once, pink). There are *p* (x-axis) memories in total, half in either category. The black dashed line indicates the initial value of the overlap (i.e., *f*).

The apparent exception is, perhaps surprisingly, when the pool of items is very small, *K* = 50 – for those it appears that the network remains highly correlated with the cue, hence with the memory itself, all the way to high values of *p*. This is due to two effects, as clarified by Fig. 2b. First, we can imagine the network as moving on a free-energy landscape (or its generalizations, the details are beyond our purposes here); for movement to be unimpeded, the landscape has to be smooth, which it is not for *K* = 50, due to the limited number of item representations dotting it. Now, the full cue does not really test the retrieval or pattern completion ability of the recurrent Potts network, but only its reluctance to drift away from the initial configuration of activity, already specified by the cue – and with a rough landscape the network is very reluctant, as it cannot effectively move. When using a partial cue, instead, e.g. *f* = 0.5, the other overlap curves do not change much, but the one for *K* = 50 starts to drop already for *p >* 50. Second, if the cue maintains nevertheless activated the items it is applied to (3 out of 5, for *f* = 0.5), there is a substantial chance, if the pool is small, that also some of the remaining items will be those appearing in the memory to be retrieved. So we can consider the small *K* case as essentially an artifact, in any case irrelevant to human memory, which represents more than 50 items.

For larger *K* values on the other hand the constraint is real, and it can be understood to a first approximation by considering the individual items as robust blocs of units that can be reactivated coherently, while the challenge for the network lies in using the item pairings, stored in the connectivity, in order to retrieve the correct configuration of *Z* items. The challenge is tougher the more memories are stored, because more pairs of items will have been stored in the connections between the Potts units. Resorting to an argument developed many years ago for the Willshaw model [23, 24], we can estimate the probability that a pair of items has *not* been stored as the probability that it has not been stored in one memory, to the power *p*: Prob(*F*_*ρσ*_ = 0) = [1 − (*Z/K*)^2^]^*p*^ *≈* exp(−*pZ*^2^*/K*^2^). If this probability becomes small, most item pairs will be encoded in the network, that will find it difficult to select those in the compositional memory. This interference effect is reduced for large *K*, but then a complementary negative effect sets in, that the network is overloaded with items. Simulations show that the two effects complement each other and lead, irrespective of the *K* value, to an effective capacity much reduced with respect to that of unitary memories.

### Memories composed with frequent and with rare items

Fig. 2c shows that the capacity constraints is almost as stringent also in a network that has stored compositional memories composed of frequent (hence, repeated) items, and *other* memories composed of rate items (in our model, appearing only once, hence unambiguously individuating the episodic memory that includes them). The effective storage capacity for the latter is a bit higher, as the signal that leads from a partial cue to reactivate the complete configuration of items is clearer, but since the noise is contributed and felt by both frequent and rare memories once they share the same network, the difference is small. Note that in Fig. 2c frequent items, from a pool of *K* = 100, are repeated as many times as those with *K* = 200 in Fig. 2b, as they appear in half of the *p* memories.

We have also simulated a network storing half compositional and half unitary memories. Unitary memories can also be conceived as composed of items appearing only once; the difference with the case above is in the learning rule, which in the compositional case of Eq. (7) assigns more weight to the individual items, because of the prefactor 1*/a*^*′*^. Overall, however, the interference resulting from the storage of the other memories is similar, and so is the resulting storage capacity for compositional and unitary memories (not shown). Note that if the latter were alone, many more of them could be stored, but since they share the connection space with compositional memories, their effective “storage capacity” is almost the same as that of compositional memories.

### Scale-free memories

Next, we consider a more realistic case in which memories include items of different frequencies. We proceed as follows: we group items into *B* bins, indexed by 1, …, *l*, …, *B*, and each bin includes *l* items (Fig. 3a). Then a memory is assembled by combining *Z* items obtained by sampling bins evenly. This results in the few items in the first bins being picked up more frequently than the many in the later bins and, as one can easily show, in an approximately *scale-free* distribution of items across memories (here, scale refers to frequency; see also the Appendix).

**Figure 3.**
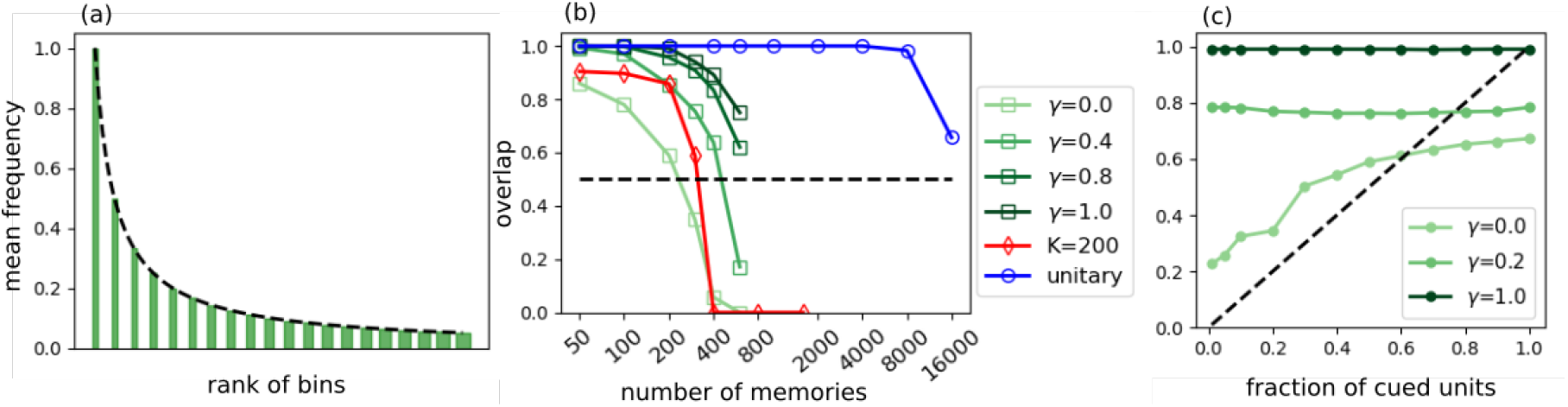
Scale-free distribution of item frequency. **(a)**: An example of distribution of item frequency with *B* = 20 bins. Bins are arranged according to the frequency of items they include along the x-axis, with frequency indicated by bin height, while bin width alludes to the number of items per bin. **(b)**: Retrieval for memories comprised of items following the frequency distribution given in (a). Colour encodes *γ* values. *f* = 0.5, *B* = 20. The red curve is for single frequency items, as in Fig. 2b. **(c)**: Similar to (b), the overlap is shown as a function of *f* for *p* = 200.

Fig. 3b (lightest green curve) shows that diversity in the distribution of item frequency has an adverse effect on storage capacity. A suitable comparison is between a scale-free distribution of items in *B* = 20 bins, which implies *B*(*B* + 1)*/*2 = 210 items overall, the lightest green curve, and compositional memories with fixed-frequency items drawn from a pool of *K* = 200 (the red curve). The comparison indicates that the more realistic, mixed distribution of item frequencies, coexisting within the same memories, does not solve the capacity constraint imposed on compositional memories; if anything, it makes it somewhat worse.

### Hippocampal inputs

The results above indicate that memory retrieval triggered by partial cues is inherently less effective with compositional memories, in which the component items have been stored on their own, than with unitary representations. This suggests that a more effective retrieval operation could be initiated by a full cue, possibly weak but full, that is, distributed over all the component items. Such a cue could come from an auxiliary compressed representation of the full memory, of the type that the hippocampus has been widely thought to store and retrieve, in turn, from partial cue.

To explore this hypothetical mechanism, we add a model hippocampal input to the compositional representation in the extended cortical network; following [22], this is simply a sustained external contribution to the signal aligning each Potts units towards the activation state it has in the memory to be retrieved. It is parametrized by a variable *γ*. In formulas, using Kronecker’s *δ* we write

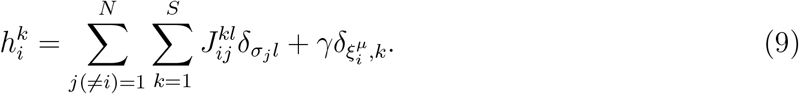

Obviously, when the factor *γ* is large enough, successful retrieval is expected to be merely transferred from the hippocampus to the neocortex, in this model, with the latter not performing any significant role. As shown in Fig. 3b, however, simulation results are complex. On the one hand, the sustained input enhances network capacity, the more the stronger it is, but without really removing the capacity limit for compositional memories, indicated by the drop in all green curves at *p* = 600. On the other hand, also a weak hippocampal input produces a noticeable effect, when the fraction of the standard partial cue is *f* = 0.5. When *p* = 200, Fig. 3c shows that even a weak sustained input, *γ* = 0.2, leads to retrieval to a level midway to that obtained with *γ* = 1.0, and as a function of *f* the same level is reached in the entire range *< f <* 1.0: in practice the hippocampal input requires only a minimal additional cue – and also when this is absent (*f ≃* 0.0) hippocampally-triggered retrieval is effective on its own.

### Triggering retrieval from frequent or from rare items

Given the interference caused by the multiple pairings of frequent items with others, in retrieving compositional memories, one may wonder whether the operation is more effective if triggered by the reactivation of the rarer items. This can be examined in the model simply by applying the partial cue *f* to the Potts unis active in the representation of the rare vs. the frequent items. In Fig. 4, this is done considering model scale-invariant representations produced with *B* = 20 bins, and applying the cue at either end of the frequency spectrum (solid curves); or by selecting memories with at least one item from the first two (frequent) or the last two (rare) bins, and averaging only over either restricted subset (dashed lines). Fig. 4a shows that without the model hippocampal input, *γ* = 0.0, there is a marked effect of where the cue is applied, but only for *f* ≤ 0.2, i.e., effectively when a single item is cued. In that case cuing a frequent item is ineffective, while cuing a rare one is (partially) effective, although the correlation with the full memory is still far from ideal (overlap just above 0.4, or 2/5 items retrieved). With weak hippocampal input, *γ* = 0.2, retrieval is still incomplete, but the effect of where the partial cue was applied is virtually erased.

**Figure 4.**
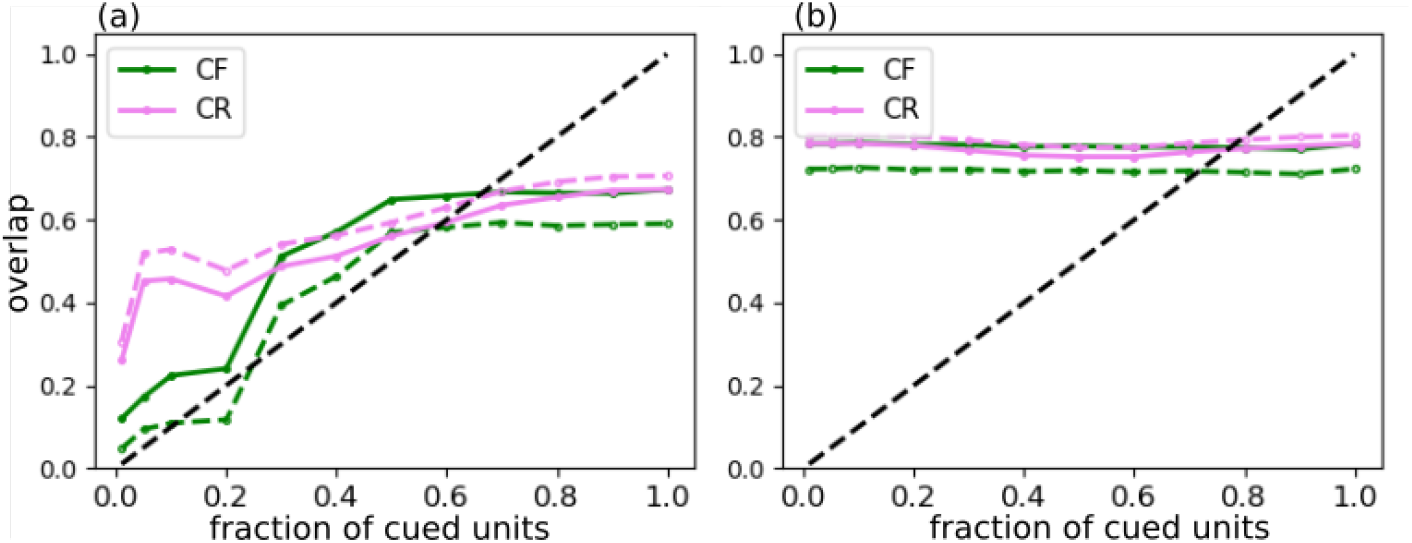
Effect of item frequency on retrieval. **(a)**: Two methods of cuing the network with a partial version of memory patterns are compared by different colours. A fraction *f* of units among *Na* active units for a memory pattern is chosen preferentially from frequent items (green) or from rare items (violet). *p* = 200, *γ* = 0.0, *B* = 20. **(b)**: Same as (a) but for *γ* = 0.2. Dashed curves show results that include only a subset of memories that include very rare or very frequent items (from the last or first two bins in the distribution).

## 4 Discussion

In this study we look at purely computational constraints for the retrieval of episodic, compositional memories, which turn out to be relatively complex to analyze, despite the artificial simplicity of the assumptions in-built in the model we have considered. To assess the range of validity of the results obtained, it is therefore useful to review the main assumptions:

- compositional episodic memories are conceived statistically as being structured in terms of items, independently drawn from a pool of such items, with no further substructure. For example, the image of a groom can be composed with that of a thick wood as well as with that of a lawn, even though weddings more often take place on the latter. In further work we shall relax this assumption by introducing structured schemata into the model.
- two distinct modes of content-addressing an episodic memory are envisaged. In the first, a partial cue sets in the active state the Potts units relative to a fraction *f* of the items composing the memory – which is intended to correspond to the initial alignment of some patches of cortex along the local attractors which represent those items, while the rest of the cortex is not aligned to anything.
- in the second mode, the hippocampus provides a sustained cue of possibly limited strength, but delivered to all relevant patches of cortex – therefore, a hippocampal index in Teyler and DiScenna’s sense [25] rather than a partial cue.
- the hippocampal representation of a compositional episodic memory, if it exists, is assumed to be unitary and not compositional, hence unrelated to the detailed semantic content of each item.
- other simplifying assumptions are more “technical”, as they relate to the Potts neural network model.

Obviously all such assumptions are extreme, and relaxing them results in some form of interpolation. This can be regarded as a general limitation of an approach which, in the trade-off between clarity and plausibility, favors the former.

### Compositionality effectively shrinks the cortex

The model offers a number of theoretical insights. One of the main findings is that the storage capacity that had been previously calculated for unitary representations [16, 21] is much higher and essentially irrelevant to that for compositional representations. The storage capacity for compositional representations is indeed constrained by factors that should be investigated further: the statistics of compositionality, the (long-range) connectivity, the plasticity that underlies the acquisition of compositional memories (expressed in the model by the “learning rule” adopted). The key finding, that is, the low storage capacity for compositional representations, may seem counter-intuitive: using a representation preassembled in blocks of units – the items – makes recall more difficult instead of facilitating it. The computational reason is that associative retrieval, in general, is robust to the interference of other memories if these produce uncorrelated fluctuations (i.e., the *noise*, in a signal-to-noise analysis) over many units or many small groups of units. If the fluctuations are coherent over large chunks of cortex, because they represent interfering items, the noise does not average out so well. It is as if compositionality nullified the key advantage of the cortex for memory – its sheer size – by obstructing the approach to the “law of large numbers”, i.e. the mutual cancellation of random fluctuations, which is key to associative retrieval. The pre-assemblage effectively reduces the size of the network to *Z*, the (average) number of items in a compositional memory; of course, only from the point of view of associative retrieval (in other respects, e.g. for representational capacity, the cortex remains huge).

### Without the hippocampus, rare elements facilitate recall

Rare elements are those shared between relatively fewer memories. The effect demonstrated in Fig. 4 reflects indeed the lower confusion associated with the retrieval cue coming from those items – they have established fewer strengthened connections to other items, and therefore are less likely to trigger the retrieval of multiple compatible memories. With the parameters we have adopted, the effect is not huge and limited to very partial cues (small *f*). Analyzing how it may scale up when cues are more detailed and the network more closely simulates a human cortex is beyond the scope of this work. We note for now that this effect, the advantage of cueing rare elements, vanishes once the hippocampus, in our model, provides a sustained full cue, even if weak, suggesting that the contribution of the hippocampus is vital to retrieve compositional memories involving highly frequent items.

### The hippocampus helps, but by brute force

A final remark on the results is that Fig. 3b indicates that the model hippocampal input does not really solve the low capacity problem. Whatever its strength *γ*, retrieval quality begins to decline at about the same memory load *p*. What happens is that in our model the hippocampus effectively takes over the retrieval task, and can send to the cortex a strong signal with its outcome, that the cortex would have been unable to get at on its own. Investigating a more significant cortical contribution, in this computational framework, probably requires a more articulated model, that we intend to analyse in future work.

### Implications for empirical research

The model can be related to a body of nascent theoretical notions and empirical data that seek to dissect the contribution of distinct brain structures to imaginative acts such as event (re)construction and mind-wandering [6]. One hypothesis is that vmPFC mediates schema-related relations among the objects in a scene, whereas the hippocampus assembles cohesive scenes [6, 26, 27]. This is consistent with the evidence that constructed experience in patients with hippocampal lesions is rich in content but lacks spatial cohesiveness, whereas that of vmPFC patients also lacks (schema-based) constitutive elements [28], and that mind-wandering is of poor episodic quality in hippocampal patients [6] vs. severely reduced in vmPFC patients [5].

In analyzing the division of labor between vmPFC and the hippocampus, a distinction that may turn out to be useful is the one analysed recently by Mullally and Maguire and involving ‘Space Defining (SD)’ and ‘Space Ambiguous (SA) objects [29, 30]. Mullaly and Maguire have suggested and shown empirically that SD objects promptly evoke a strong sense of a surrounding 3D space. An example SD object is a couch, which promptly evokes a sense of a surrounding 3D space compatible with a living room and not with other types of spatial layouts; SD objects define (identify) the space they fit in. A fly, an example SA object, does not. SA objects are compatible with, and shared between, many spatial layouts. Consistent with the prominent role of space processing for mental construction, SD objects are preferentially chosen as the initial building block to mentally construct a scene, and are picked last to be removed from a mental scene [30]. Processing of SD and SA stimuli is associated with different activity in the parahippocampal cortex [29], the superior temporal gyrus, and vmPFC [31], in line with the different functional properties of the two classes of items.

The SD-SA distinction must be considered together (and not confounded) with another independent distinction, that between objects that are more or less likely to be associated with other objects or related concepts [32], and hence trigger their activation [29]. Although SD objects tend to be evocative of content (associated with other objects/concepts), as in the previous example of the couch, which can easily activate, in addition to the 3D space of a living room, the image of a nearby coffee table or TV, the SD/SA and contextual richness dimensions are distinct, and dissociable from the one another both behaviorally and neurally [29]. We have recently isolated, for example, SD objects high in contextual associations (eg: swing), SD objects low in contextual associations (eg: chair), SA objects high in contextual associations (eg: fishing rob), and SA objects low in contextual associations (eg: belt) to be used as cues for event construction (Stendardi *et al*., in preparation).

In the present model, rare items can be taken to more immediately evoke a constellation of other items, because, being rare, they have been associated strongly with a small number of other items and contexts. This is the case for SD items, especially those with low levels of contextual associations, which evoke virtually unique contexts. Cueing a rare (e.g., SD) item is likely more effective in triggering memory retrieval, as competition between memories sharing that item is less likely. Our computational findings indicate that if and only if the hippocampal input is damaged or reduced, a partial cue applied to a rare item is more effective in triggering accurate memory retrieval than one applied to a frequent item. It would be interesting to investigate, therefore, if the advantage in event construction observed for SD vs. SA items is more pronounced in the case of reduced input from the hippocampus, for example testing patients with hippocampal damage or using tasks that make heavier demands on neocortical regions vs. the hippocampus (e.g., priming).

As of now, it remains unclear to what extent the model captures the *spatial* nature of memories for multiple items in visual scenes (which is integral to the SD/SA distinction); but it is clear that it fails to consider more structured constructs, usually referred to as *schemata*. These can be elaborated in at least two different dimensions. One is to consider schemata as groups of items that often occur together as components of wider compositional scenes, irrespective of exact timing relations. For example, the memory of a countryside wedding may include a makeshift religious ceremony and a festive meal, each of which includes distinct items bound together into those two schemata. A second dimension is the temporal one. If two items A and B when they co-occur do so in a fixed succession, such as the discussion of the Thesis and the friends’ congratulations, proper recall would entail reactivating their representation in the same order. Ultimately, along both dimensions one moves away from the snapshot character of simple episodic memories, taking some steps towards their *semantization*.

Developing our current computational model along the first dimension involves considering some form of nested probability distributions, which opens up a very large space of possibilities, so that it is probably wise to focus on a specific set of empirical data. Along the second dimension, instead, there is a straightforward neural mechanisms that favors the ordered reactivation of the representations of two items A and B: to enhance the connections from the units active in A to those active in B, and not viceversa. If a spatial relation is captured, in part, by the availability of both options, scanning A→B as well as B→A, a temporal relation singles out A→B. Correctly reactivating all the temporal relations in an episode that has been experienced could be challenging for the cortex, but a partial reactivation that follows several originally distinct paths may in fact be the substrate for the generative process envisaged by Barry and Maguire [8].

Ongoing work is exploring whether assigning such temporal structure to a simple composition of items can indeed facilitate their ordered reactivation, irrespective of the model hippocampal signal. If so, the model may be useful to better understand the phenomenology presented by vmPFC patients [9] and also, perhaps, the third and most sophisticated form of the imaginative faculty, as conceived by Dante.

## Acknowledgments

We are grateful to Andrea Tabarroni, professor of Medieval Philosophy, who showed us the passage from Dante’s Comedy on the imaginative faculty.

## Author contributions

The model was developed by KIR and AT based on ongoing empirical research by DS and EC. Simulations were run by KIR, and the results discussed by all coauthors, who together wrote the paper.

Appendix

## Making memory representations

We construct representations of compositional memories in two steps. In the first step, we assign *Z* items to each memory. This is done either by sampling items evenly, so that on average they all occur with the same frequency, or unevenly, as described in the text, for example with the quasi-scale-free procedure discussed below, and represented in Fig. 3a. In the second step, we write a representation of each memory by merging representations of its *Z* items. The only issue in doing so is that there are some units that are shared by more than one item. This would lead to representations with sparsity (fraction of active units) less than *a*. In order to constrain all memories to have the same sparsity *a*, we compute the “fields” 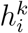 of all units, by assuming that the *Z* items of a particular memory are activated. Then we select the *Na* units (and their states) which receive the largest field, to define them as the representation of this particular memory.

### Scale-free item frequency

Scale-free distributions have been invoked as a simple description of many natural phenomena, and there is considerable controversy as to the ideas that have been put forward [33, 34]. There has been also considerable work on the scale-invariant distribution of objects of different sizes in natural scenes, which is closer to being relevant for the compositionality of memory for scenes [35]. Here our intent is merely practical, however: to generate a simple distribution of frequencies, which does not involve an extra arbitrary parameter. The distribution described in the text is approximately scale free, because no such parameter is introduced explicitly, although implicitly the number *B* of bins sets the upper and lower ends of the frequency range with which items are assigned to memories: from about *pZ/B* to *pZ/B*^2^ times. Within this range, each “frequency scale” is approximately represented evenly.

## Default parameters

When not varied systematically, parameters of the Potts model are set as Table 1.

